# Hibernation reduces GABA signaling in the brainstem to enhance motor activity of breathing at cool temperatures

**DOI:** 10.1101/2023.10.09.561534

**Authors:** Sandy E. Saunders, Joseph M. Santin

**Affiliations:** University of Missouri-Columbia, Division of Biological Sciences, Missouri, United States of America

## Abstract

**Background:** Neural circuits produce reliable activity patterns despite disturbances in the environment. For this to occur, neurons elicit synaptic plasticity during perturbations. However, recent work suggests that plasticity not only regulates circuit activity during disturbances, but these modifications may also linger to stabilize circuits during future perturbations. The implementation of such a regulation scheme for real-life environmental challenges of animals remains unclear. Amphibians provide insight into this problem in a rather extreme way, as circuits that generate breathing are inactive for several months during underwater hibernation and use compensatory plasticity to promote ventilation upon emergence.

**Results:** Using *ex vivo* brainstem preparations and electrophysiology, we find that hibernation in American bullfrogs reduces GABA_A_ receptor (GABA_A_R) inhibition in respiratory rhythm generating circuits and motor neurons, consistent with a compensatory response to chronic inactivity. Although GABA_A_Rs are normally critical for breathing, baseline network output at warm temperatures was not affected. However, when assessed across a range of temperatures, hibernators with reduced GABA_A_R signaling had greater activity at cooler temperatures, enhancing respiratory motor output under conditions that otherwise strongly depress breathing.

**Conclusions:** Hibernation reduces GABA_A_R signaling to promote robust respiratory output only at cooler temperatures. Although animals do not ventilate lungs during hibernation, we suggest this would be beneficial for stabilizing breathing when the animal passes through a large temperature range during emergence in the spring. More broadly, these results demonstrate that compensatory synaptic plasticity can increase the operating range of circuits in harsh environments, thereby promoting adaptive behavior in conditions that suppress activity.

## Background

Animals have the mysterious ability to produce reliable behaviors while navigating environments that should otherwise disrupt activity of the nervous system [1]. This is thought to occur, in part, through a set of compensatory mechanisms that sense activity perturbations, and in turn, adjust neuronal properties to counteract the disturbance. For example, if activity falls, neurons increase synaptic excitation, decrease inhibition, and alter ion channel densities to enhance neuronal excitability [2–5]. This framework, termed “homeostatic plasticity,” has been foundational for the past 30 years [6], but it has been challenging to link ethologically relevant behaviors in animals, as neurons often need to be perturbed artificially or pathologically to reveal homeostatic responses [7–12].

Certain organisms inhabit natural environments with potent abiotic stressors, such as pH, temperature, and hypoxia, which can cause activity challenges within the nervous system [13]. An extreme example of this problem is encountered by hibernating frogs. Like most vertebrate species, rhythmic neural circuits in the brainstem generate breathing to meet metabolic demands of the organism. However, in the cold hibernation environment frogs may spend several months underwater using only skin for gas exchange [14], while respiratory motor behavior is completely suspended [15]. To successfully emerge following months underwater, frogs employ various compensatory mechanisms to maintain these networks so that they can work effectively when needed. This includes a classic activity-dependent mechanism of compensatory plasticity called “synaptic scaling” at excitatory synapses [16, 17], and also metabolic adjustments that improve performance in hypoxia [18, 19]. As amphibians use synaptic plasticity to overcome this large environmental challenge, they provide an intriguing opportunity to understand how plasticity mechanisms are integrated within animals to promote adaptive behavior.

Compensatory forms of synaptic plasticity are often interpreted as a conceptually simple regulatory regime used to counteract activity disturbances (e.g., when activity goes down, excitatory synaptic strength goes up). However, experimental and modeling work indicates that when circuits are disrupted, compensatory adjustments can also “prime” circuits to perform better during future perturbations and influence the capacity for subsequent plasticity [20–22]. Thus, we hypothesized that the hibernation environment not only triggers synaptic plasticity, but also that it induces network modifications to expand the operating range in challenging environments encountered following the initial activity challenge. To test this in an ethological context, we built upon previous work demonstrating that hibernation drives apparent homeostatic plasticity at excitatory synapses [16, 17] and addressed whether inhibitory synapses contribute as well. We then asked how plasticity through inhibition influences motor function of breathing at colder temperatures, where neuronal activity is otherwise markedly suppressed. Here, we report that hibernation strongly reduced GABAergic inhibition, consistent with a classic response to chronic network inactivity [3, 23]. Despite the critical role of inhibition in this network [24], circuit function appeared surprisingly normal at warm temperatures. However, when we assessed activity across a range of temperatures, GABA_A_R plasticity served mainly to offset the depressive effects of acute cooling, boosting the strength and frequency of motor output at lower temperatures. Given that these animals must pass through a range of temperatures as they reestablish life on land after hibernation [25, 26], network modifications that improve network activity within this range seem to play an adaptive role. Therefore, although GABAergic plasticity has no obvious impact on baseline motor function, it expands the operating range of the respiratory circuit in an otherwise suppressive environment to promote adaptive behavior.

## Results

Respiratory motor output in adult frogs is thought to arise *via* central pattern-generating mechanisms consistent with reciprocal inhibition involving GABA [24].

Rhythmic activity in these premotor neurons is then carried to motor pools through interneuronal pathways that control motor outflow to respiratory muscles [27]. Activity of this network can be assessed in isolated brainstems *in vitro* which produce rhythmic motor output on a variety of cranial nerves including the trigeminal, vagal, and hypoglossal nerve root. Although the activity pattern of these nerves may differ, the output produced is largely synchronous providing a representation of the motor output that resembles lung ventilation in the whole animal [28, 29]. Here, we measured activity of the vagus nerve to monitor respiratory-related motor output from the intact brainstem preparation (cranial nerve X; CNX; schematic in Figure 1).

**Fig. 1.**
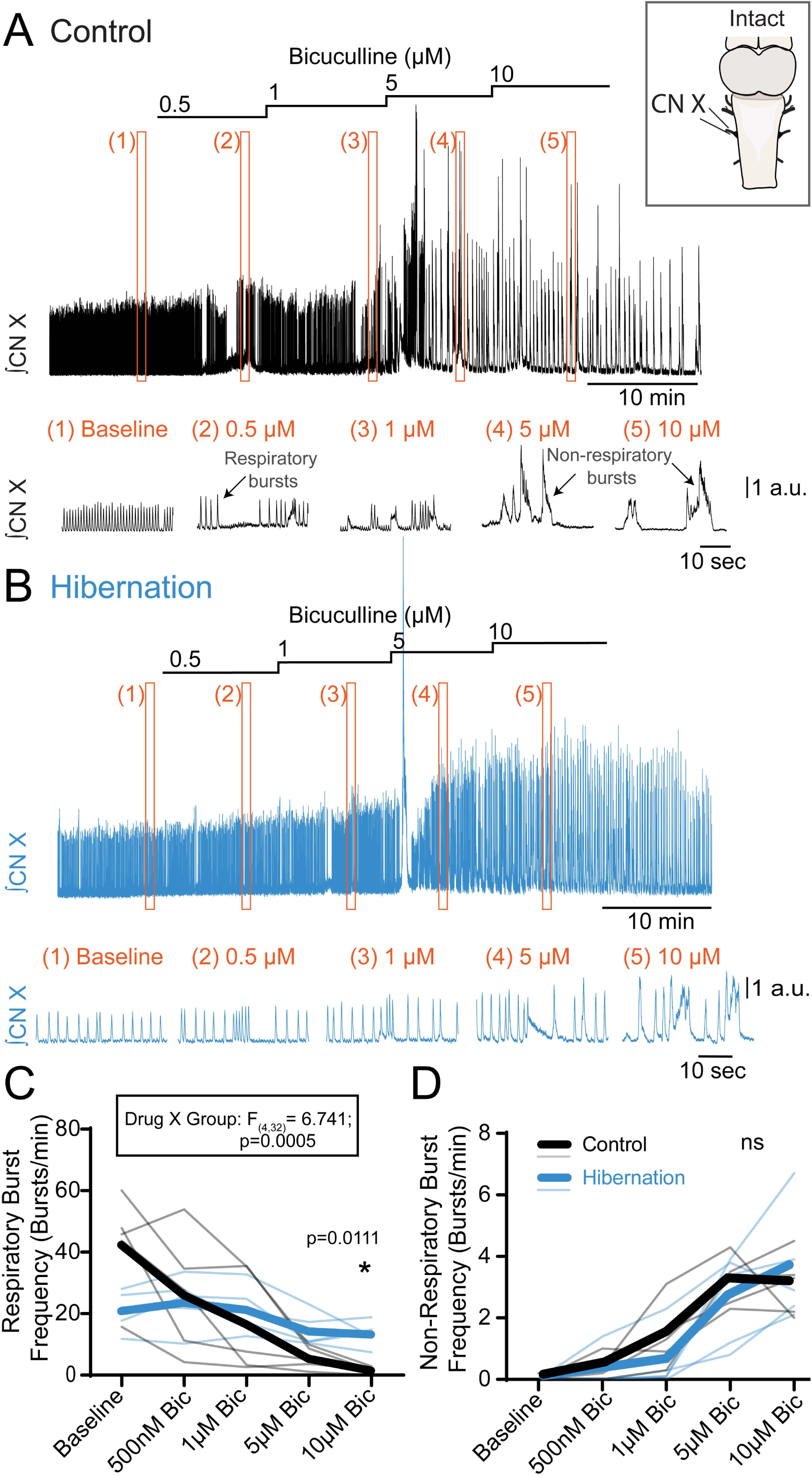
GABAergic tone that controls respiratory burst frequency strongly decreases following hibernation. Integrated vagus nerve activity (CN X) was rhythmic in fully intact brainstem preparations (top right box) from control and hibernated animals. A) Top, continuous recording of CNX during perfusion of increasing doses of bicuculline in a control preparation (black). Bottom, zoomed in view of epochs (orange boxes) indicated in continuous trace directly above. In control preparations, bath application of bicuculline led to a dose dependent decrease in respiratory burst frequency and dose-dependent increase in non-respiratory burst frequency. B) Top, continuous recording of CNX during perfusion of increasing doses of bicuculline in a preparation from a hibernated animal (blue). Bottom, zoomed in view of epochs (orange boxes) indicated in continuous trace above. Respiratory output from hibernated preparations was insensitive to systemic application of bicuculline, however non-respiratory motor bursting increased in a dose-dependent manner like controls. C) Summary of bicuculline-mediated changes in respiratory burst frequency. There was a significant interaction between drug and group by two-way ANOVA (F_(4,32)_=6.741; p=0.0005). Additionally, at 10 µM bicuculline, respiratory burst frequency from controls was significantly lower than hibernated preparations by Holm-Sidak’s post-hoc test (p=0.0111; n=5/group). D) Summary of bicuculline-mediated changes in non-respiratory burst frequency. Bicuculline mediated similar increases in non-respiratory bursting in control and hibernated groups. There was no significant interaction between drug and group by two-way ANOVA (F_(4,32)_=0.7948; p=0.5373; n=5/group). a.u., arbitrary units. In summary plots, thick lines represent the group average and thin lines are responses of individual preparations.

To investigate plasticity in GABAergic control of the respiratory network, we first bath-applied bicuculline, a GABA_A_ receptor antagonist, to intact brainstem preparations. Consistent with an important role for GABA_A_Rs in rhythmic motor activity, bicuculline caused a dose-dependent decrease in burst frequency in control brainstems at room temperature (Baseline, 42.60 ± 16.32 bursts/min; 0.5 µM Bicuculline, 26.32 ± 19.68 bursts/min; 1 µM Bicuculline, 16.86 ± 17.03 bursts/min; 5 µM Bicuculline, 5.66 ± 3.61 bursts/min; n=5, Figure 1A,C), with most having little-to-no activity at the highest dose (10 µM Bicuculline, 1.74 ± 1.10 bursts/min; n=5, Figure 1C). Under the same experimental conditions, hibernators strongly resisted bicuculline (Baseline, 21.12 ± 6.53 bursts/min; 0.5 µM Bicuculline, 23.80 ± 8.72 bursts/min; 1 µM Bicuculline, 21.52 ± 7.71 bursts/min; 5 µM Bicuculline, 14.50 ± 5.54 bursts/min; n=5, Figure 1B,C), with respiratory activity persisting at the highest dose (10 µM Bicuculline, 13.50 ± 4.12 bursts/min; n=5, Figure 1B,C). In regard to motor amplitude, two-way ANOVA revealed a significant main effect of bicuculline on normalized motor amplitude (F_(2.055, 16.44)_ = 4.499, P=0.0269) but not a significant effect of group or an interaction between bicuculline and group as we saw for frequency. However, this bicuculine effect is likely to be minor, as post hoc analysis showed that significantly larger bursts compared to baseline occurred at 5 μM in controls (p=0.010; Holm-Sidak Multiple Comparison’s test), while all other comparisons to baseline were not significantly different from baseline.

Loss of GABA_A_R signaling appeared to be localized to breathing circuits. In both control and hibernators, non-respiratory motor activity (long duration bursts with qualitatively different shape than respiratory bursts [30, 31]) were rare at baseline with only 1 control preparation containing these bursts. Interestingly, and despite a clear group x bicuculine interaction for respiratory burst frequency, we did not observe this same response for non-respiratory bursts (group x bicuculline interaction in two-way ANOVA; p=0.5373). Non-respiratory bursts emerged similarly in a manner that depended on the dose of bicuculline (Figure 1A-B,D) in both controls (Baseline, 0.04 ± 0.09 bursts/min; 0.5 µM Bicuculline, 0.44 ± 0.39 bursts/min; 1 µM Bicuculline, 1.44 ± 1.05 bursts/min; 5 µM Bicuculline, 3.18 ± 0.81 bursts/min; 10 µM Bicuculline, 3.08 ± 1.01 bursts/min, n=5) and hibernators (Baseline, 0.00 ± 0.00 bursts/min; 0.5 µM Bicuculline, 0.28 ± 0.63 bursts/min; 1 µM Bicuculline, 0.56 ± 0.98 bursts/min; 5 µM Bicuculline, 2.62 ± 1.50 bursts/min; 10 µM Bicuculline, 3.60 ± 1.86 bursts/min, n=5).

Given the difference in how respiratory and non-respiratory bursts alter sensitivity to bicuculline, hibernation strongly reduces the role of GABA_A_Rs in a way that appears specific to circuits that generate breathing but not non-respiratory behaviors.

The previous experiment demonstrated that GABA_A_Rs have a strongly reduced role in the production of the respiratory rhythm after hibernation. Next, we assessed whether plasticity influenced the capacity of the respiratory network to be modulated by GABA_A_R activation. Therefore, we bath-applied muscimol, a GABA_A_R agonist, to globally activate all GABA_A_Rs that influence activity. In controls, the lowest dose of muscimol (500 nM) did not alter respiratory burst frequency (Baseline, 8.27 ± 3.7 bursts/min *vs.* 500 nM muscimol, 7.20 ± 4.0 bursts/min; n=5). Raising the concentration to 1 µM caused a significant decline (1 µM muscimol, 3.47 ± 1.9 bursts/min; p= 0.0282; n=5; Figure 2A,C), and exposure to 3 μM silenced all preparations (5 out of 5). In contrast, preparations from hibernators did not significantly alter respiratory burst frequency between baseline and any dose of muscimol (Baseline, 11.00 ± 2.1 bursts/min; 500 nM muscimol, 11.93 ± 1.8 bursts/min; 1 µM muscimol, 14.47 ± 2.8 bursts/min; 3 µM muscimol, 6.47 ± 7.3 bursts/min; n=5; Figure 2B-D), with 5 out of 5 preparations maintaining respiratory bursting at the highest dose. While suppression of network activity with both the antagonist and agonist may seem paradoxical, these results are consistent with the necessity of phasic inhibition for generation of respiratory activity and neuronal silencing through further activation ofGABAergic inhibition, respectively. Indeed, in other circuits that use inhibition for rhythmogenesis, antagonism of GABA_A_Rs disrupts network dynamics required for network activity [32], while agonism of GABA_A_Rs also suppresses activity through membrane hyperpolarization [33]. With both agonist and antagonist experiments, these results show that hibernation leads to a large reduction in the ability of GABA_A_Rs to generate and modulate breathing.

**Fig. 2.**
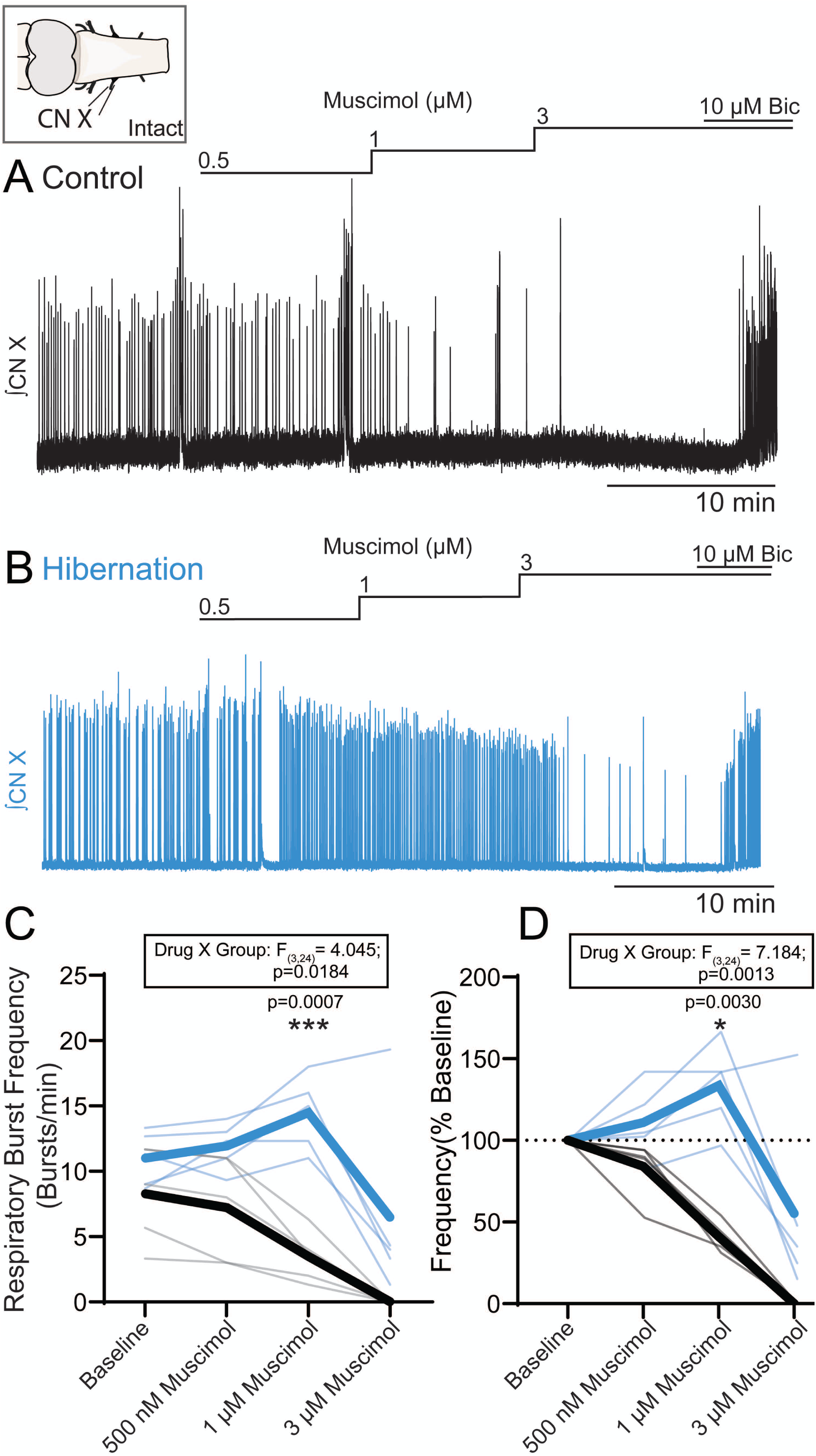
Hibernation decreases sensitivity to GABA_A_R agonist (muscimol)-mediated respiratory frequency decline. Integrated vagus nerve activity (CN X) was rhythmic in fully intact brainstem preparations (top left box) from control and hibernated animals. A) Continuous recording of CNX during perfusion of increasing doses of muscimol in a control preparation (black). In control preparations, respiratory output was unchanged by bath application of 500nM muscimol but decreased following elevation to 1µM muscimol and ultimately stopped following further elevation to 3µM muscimol. Preparations were recovered by exposure to bicuculine, demonstrating specificity of the GABA_A_ receptor agonist. B) Continuous recording of CNX during perfusion of increasing doses of muscimol in a preparation from a hibernated animal (blue). Preparations from hibernated animals were relatively insensitive to muscimol mediated changes in respiratory burst frequency compared to controls. C) Summary of muscimol-mediated changes in respiratory burst frequency. There was a significant interaction between drug and group by two-way ANOVA(F_(3,24)_=4.045; p=0.0184). Additionally, at 1 µM muscimol, respiratory burst frequency from controls was significantly lower than hibernated preparations by Holm-Šidák’s post-hoc test (p=0.0007, n=5/group). D) Summary of muscimol-mediated changes in respiratory burst frequency relative to baseline. There was a significant interaction between drug and group by two-way ANOVA(F_(3,24)_=7.184; p=0.0013). Additionally, at 1 µM muscimol, respiratory burst frequency from controls was significantly lower than hibernated preparations relative to baseline by Holm-Sidak’s post-hoc test (p=0.0030, n=5 per group). In summary plots, thick lines represent the group average and thin lines are individual preparations.

The previous experiments identify that hibernation reduces the role of GABA_A_Rs in generating and modulating the respiratory rhythm. At the level of the motoneuron, an activity-dependent mechanism known as “synaptic upscaling” strengthens excitatory synapses in response to hibernation [16, 17], aligning with classic homeostatic responses to chronic network inactivity [4]. Thus, we tested if hibernation also “downscales” postsynaptic GABA_A_Rs to reduce GABA_A_R inhibition. We used patch clamp electrophysiology to record miniature inhibitory postsynaptic currents (mIPSCs) carried by GABA_A_Rs in identified vagal motoneurons that cause breathing in amphibians. The GABA_A_R currents were isolated in TTX, strychnine (strych), and DNQX to block presynaptic action potentials, glycine receptors, and AMPA-glutamate receptors, respectively. GABA_A_R minis were recorded from a holding voltage of -60 mV. The Nernst potential for Cl^-^ under our experimental conditions was ∼5 mV; thus, inward currents in the presence of strychnine and DNQX represent minis that arise from GABA_A_Rs. This was confirmed in each experiment by applying bicuculline (bic) follow each recording. We did not observe changes in the mIPSC amplitude (p=0.5218; unpaired t test), charge transfer (p=0.4391; unpaired t test), rise time (p=0.1029; unpaired t test), frequency (p=0.5576; unpaired t test), as well as neuronal input resistance after hibernation (p=p=0.3647; unpaired t test) (Figure 3A-C). Therefore, hibernation does not influence postsynaptic GABA_A_Rs and is unlikely to account for the loss of GABA_A_R signaling at the network level.

**Fig. 3.**
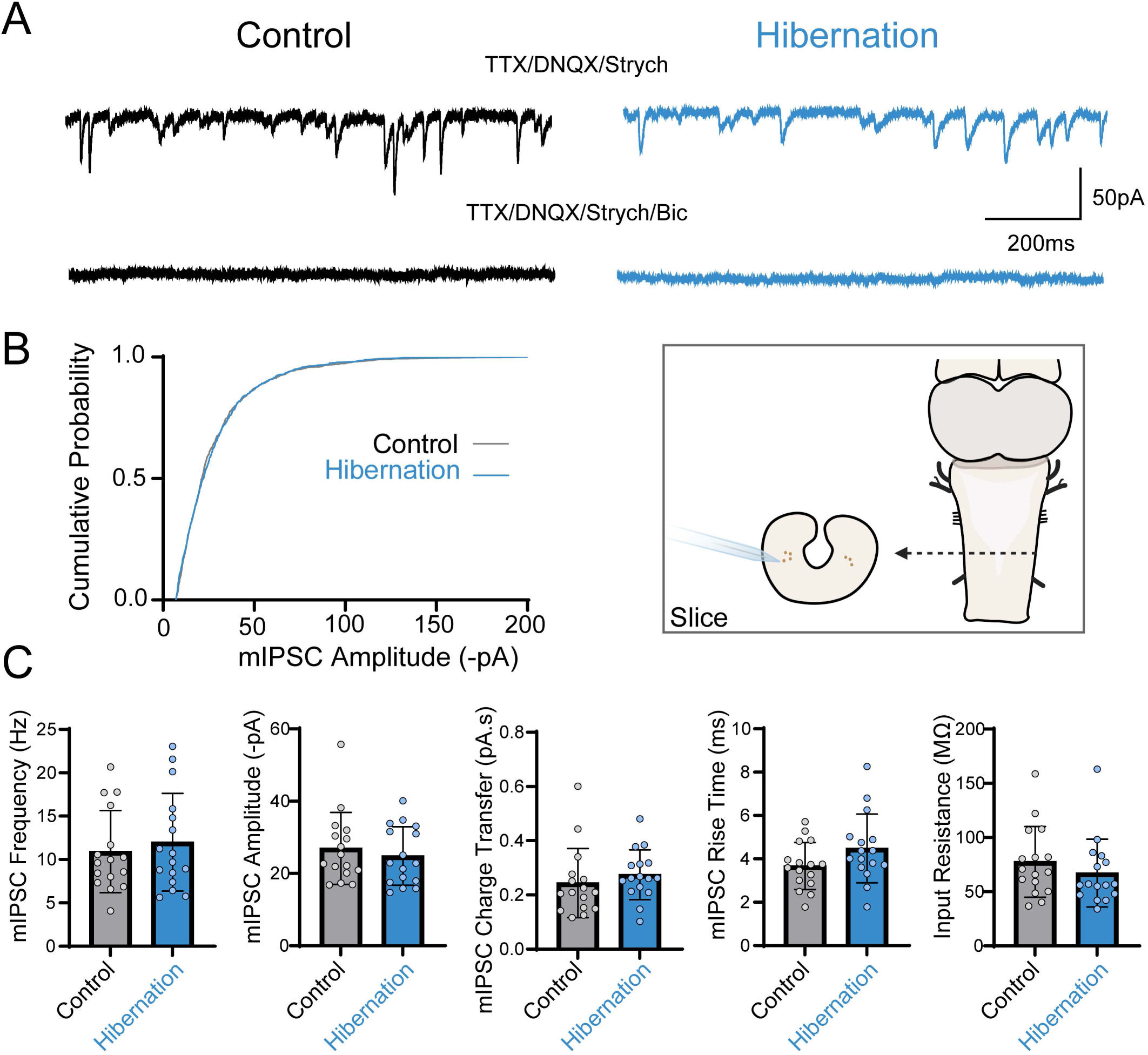
Postsynaptic GABAergic currents in motoneurons are unchanged following hibernation. A) Example voltage clamp traces of GABAergic miniature inhibitory postsynaptic currents (mIPSCs; holding potential = -60 mV; 250 nM TTX, 10 µM DNQX, 5 µM strychnine) from control (black) and hibernated (blue) identified vagal motor neurons in brain slice (middle right box). Addition of 50 µM bicuculline (Bic) to the bath eliminated mIPSCs in both groups, confirming GABAergic identity. B) Cumulative probability histograms of mIPSC amplitudes from control neurons (black line), hibernated neurons (blue line) appeared nearly identical. Accordingly, there was no statistical difference between distributions from control and hibernated cells by Kolomogorov-Smirnov test (p = 0.5041). C) Summary data. There was no statistical difference in mIPSC frequency, average amplitude, rise time, charge transfer, and neuronal input resistance by unpaired t-test (p > 0.05; n= 16 control cells; n=16 hibernated cells). Error bars are standard deviation (SD).

As the quantal GABA_A_R currents were unchanged on the postsynaptic motoneuron, we assessed if loss of the GABAergic tone is associated with reduced network-driven presynaptic input onto motoneurons, like that seen in sensory deprivation models of homeostatic plasticity [10, 11]. Specific respiratory synapses are not yet tractable for stimulation experiments in this system; therefore, we developed an approach to circumvent this issue. We used a novel semi-intact preparation that permits recording of individual motoneurons receiving excitation and inhibition from rhythmic premotor circuits that shape motoneuron firing of breathing (Figure 4A) [34]. As shown in Figure 4A, vagal motor neurons fire bursts of action potentials during respiratory population motor activity, reflecting activity of breathing at a single-cell resolution. Since the postsynaptic receptor density of GABA_A_Rs does not appear to change within these neurons after hibernation (Fig. 3), any difference in the role of GABAergic synapses for the control of motoneuron firing rate would reflect changes in network-driven presynaptic GABA release. To isolate GABA_A_Rs that control respiratory-related firing of motoneurons, we focally applied bicuculline to the motoneuron cell body and surrounding region *via* local pressure injection while recording its activity shaped by excitation and inhibition from rhythmic premotor inputs in the intact network. We simultaneously recorded rhythmic vagal motoneuron activity and the trigeminal nerve (a nerve that innerves the buccal floor to drive air into the lungs and activates near-synchronously with the vagal outflow) to monitor respiratory circuit output. We used the trigeminal nerve (cranial nerve V; CN V) in these experiments because physical constraints in the bath and the focal drug delivery system prevented simultaneous recording of the vagus nerve.

**Fig. 4.**
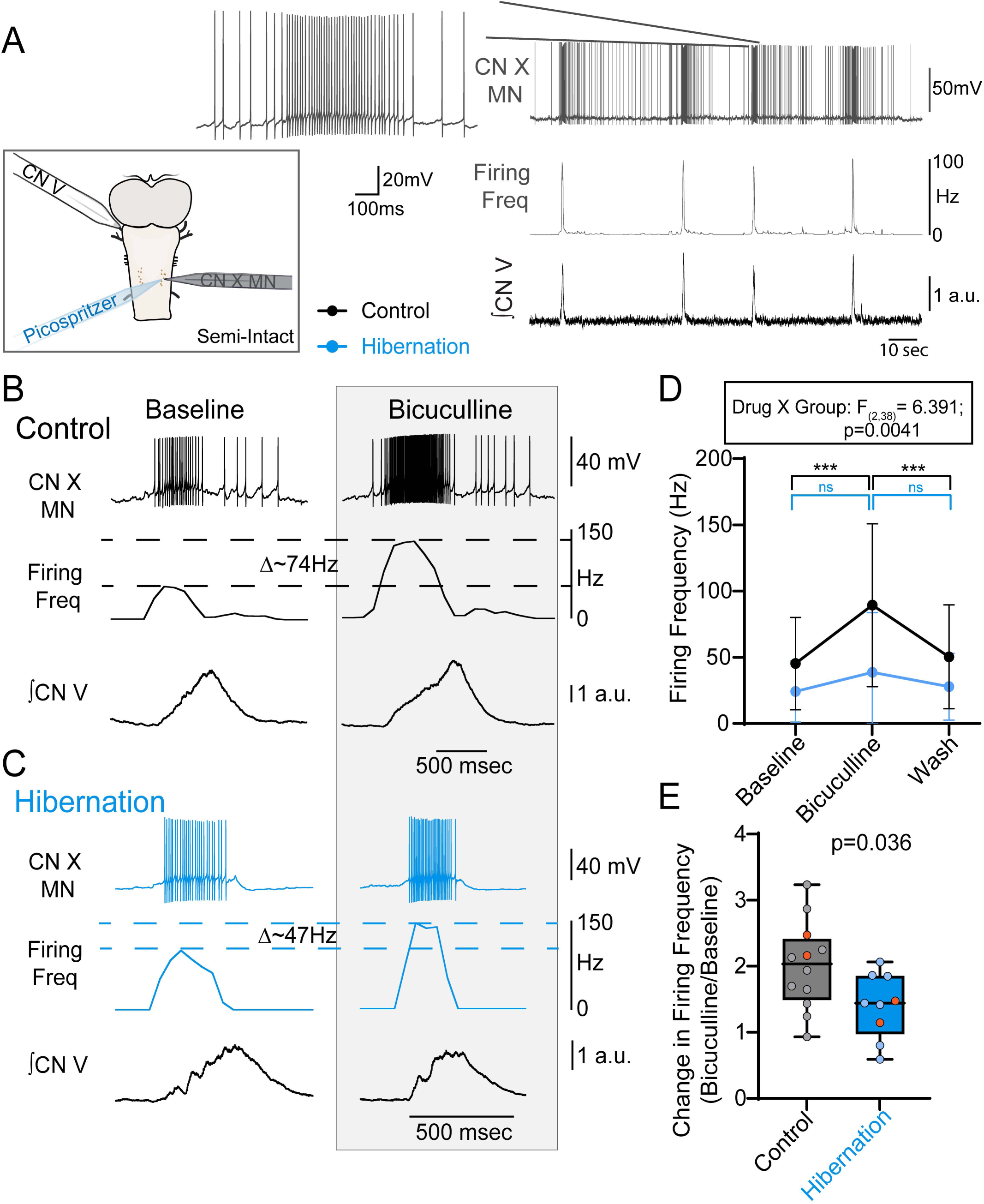
Hibernation decreases the influence of presynaptic GABA on motoneuron firing. A) Integrated network output recorded on the trigeminal nerve (CN V) and simultaneous whole-cell current clamp recordings of vagal motor neurons (CN X MN) driven by the respiratory circuit in the semi-intact brainstem preparation (top left box). A pipette was used to focally apply bicuculline onto the recorded vagal motor neuron to isolate the GABAergic tone on individual neurons. An example respiratory neuron (CN X MN, gray, top right) fired rhythmic bursts, characterized by transient increases in firing frequency (Firing Freq, light gray, middle top right) during respiratory motor output (Black, bottom top right, CN V). B) Example respiratory motor activity in a control preparation (black). Example vagal motor neuron (CN X MN) fires bursts of action potentials during the respiratory burst (CN V). Focal application of bicuculline (gray box) to the motor neuron increases neuronal burst firing frequency (Firing Freq). C) Example respiratory motor activity in a preparation from a hibernated animal (blue). Example vagal motor neuron (CN X MN) also fired bursts of action potentials during the respiratory burst (CN V). Focal application of bicuculline (gray box) to the motor neuron increased neuronal burst firing frequency (Firing Freq, Δ∼47Hz), but not to the same extent as the control neuron (Firing Freq, Δ∼74Hz). D) Summary of bicuculline-mediated increases in motor neuron firing frequency. There was a significant interaction between drug and group by two-way ANOVA (F_(2,38)_=6.391; p=0.0041). Furthermore, there was a significant increase in burst firing frequency following focal application of bicuculline in control but not hibernated cells by Holm-Sidak’s post-hoc test (***p<0.0001; n= 12 control cells; n=9 hibernated cells). E) Summary of bicuculline-mediated fold change in burst firing frequency from baseline. Control cells had a significantly larger bicuculline-mediated fold change in burst firing frequency than hibernated cells by unpaired t-test (p=0.036; n= 12 control cells; n=9 hibernated cells). Orange cells were recorded in loose-patch configuration. Box plots represent interquartile range and whiskers represent the minimum to maximum values. a.u., arbitrary units. Error bars are standard deviation (SD).

Motoneurons from both groups had similar average burst firing rates during breathing and resting membrane potentials in between respiratory bursts (firing rate: control, 45.26±34.8 Hz vs. hibernation, 24.19±23.0 Hz; p=0.133, unpaired t-test; membrane potential: control, -61.52 ± 4.6 mV vs. hibernation, -59.24 ± 6.5 mV, p=0.4094 unpaired t-test). In controls, local application of bicuculline led to a reversible increase in average firing frequency during the respiratory burst (Baseline, 45.26 ± 34.8 Hz; Bicuculline, 89.36 ± 61.4 Hz; p<0.0001; Holm-Sidak multiple comparisons test; n=12; Figure 4B,D). This was presumably not due to changes in the chloride gradient, because two neurons in the control group that were recorded in the “loose patch” configuration, which does not disturb intracellular milieu (Fig. 4E shown in orange), more than doubled their firing rates during bicuculline exposure, aligning with data from the whole-cell mode. These results demonstrate that phasic inhibition onto motoneurons dampens firing rate during the respiratory burst. In contrast, local bicuculline had no significant effect on the firing rate of motoneurons from hibernators (Baseline, 24.19 ± 23.0 Hz; Bicuculline, 38.75 ± 44.9 Hz; p=0.2629; n=9; Holm-Sidak multiple comparisons test; Figure 4C-E). Additionally, there was no change in the interburst (resting) membrane potential during focal application of bicuculline in either control (Baseline: -61.52 ± 4.6 mV; Bicuculline: -60.92 ± 4.9 mV; n=10) or hibernation groups (Baseline: -59.24 ± 6.5 mV; Bicuculline: -59.53 ± 6.5 mV; n=7), further supporting that phasic respiratory-related inhibition influences motoneuron firing rate rather than a tonic GABA_A_R current. Taken together with the mIPSC results, network-driven GABA release appears to be reduced in a way that no longer dampens the firing rate of motoneurons during activity associated with breathing.

These results demonstrate a critical role for GABA_A_Rs in generating and modulating the respiratory rhythm, as well as controlling motoneuron firing rate in control frogs. Therefore, we were surprised that baseline network frequency on average (Fig. 1C, p=0.2021) and motor burst morphology (Figure 5) in hibernators did not change despite a rather dramatic loss of GABA_A_R signaling in the network. These results mirror ventilation data *in vivo*, whereby breathing frequency and breath volume were the same after hibernation [35]. Despite these similarities at warm temperatures, frogs likely need to restart breathing to some degree at temperatures around 8°C to maintain aerobic metabolism [35, 36] and then produce reliable rhythmic output at temperatures above 13°C to maintain metabolic homeostasis as they reestablish a warmer life on land [25, 26]. Yet, cold temperatures ≤15°C strongly depress the network, making it difficult to generate respiratory activity [15, 37]. Interestingly, cooling does not suppress breathing exclusively through “passive” effects on cellular properties (*e.g*., slower action potentials, decreased rates of synaptic transmission), but rather, through noradrenergic signaling. Indeed, blocking α adrenergic receptors blunts decreases in activity during cooling, and the actions of norepinephrine on respiratory activity in the frog brainstem are known to act through GABAergic pathways [38, 39]. This led us to test whether decreased GABA_A_R signaling enhances respiratory motor output at cold temperatures faced by frogs when they need to resuscitate breathing while emerging from hibernation.

**Fig. 5.**
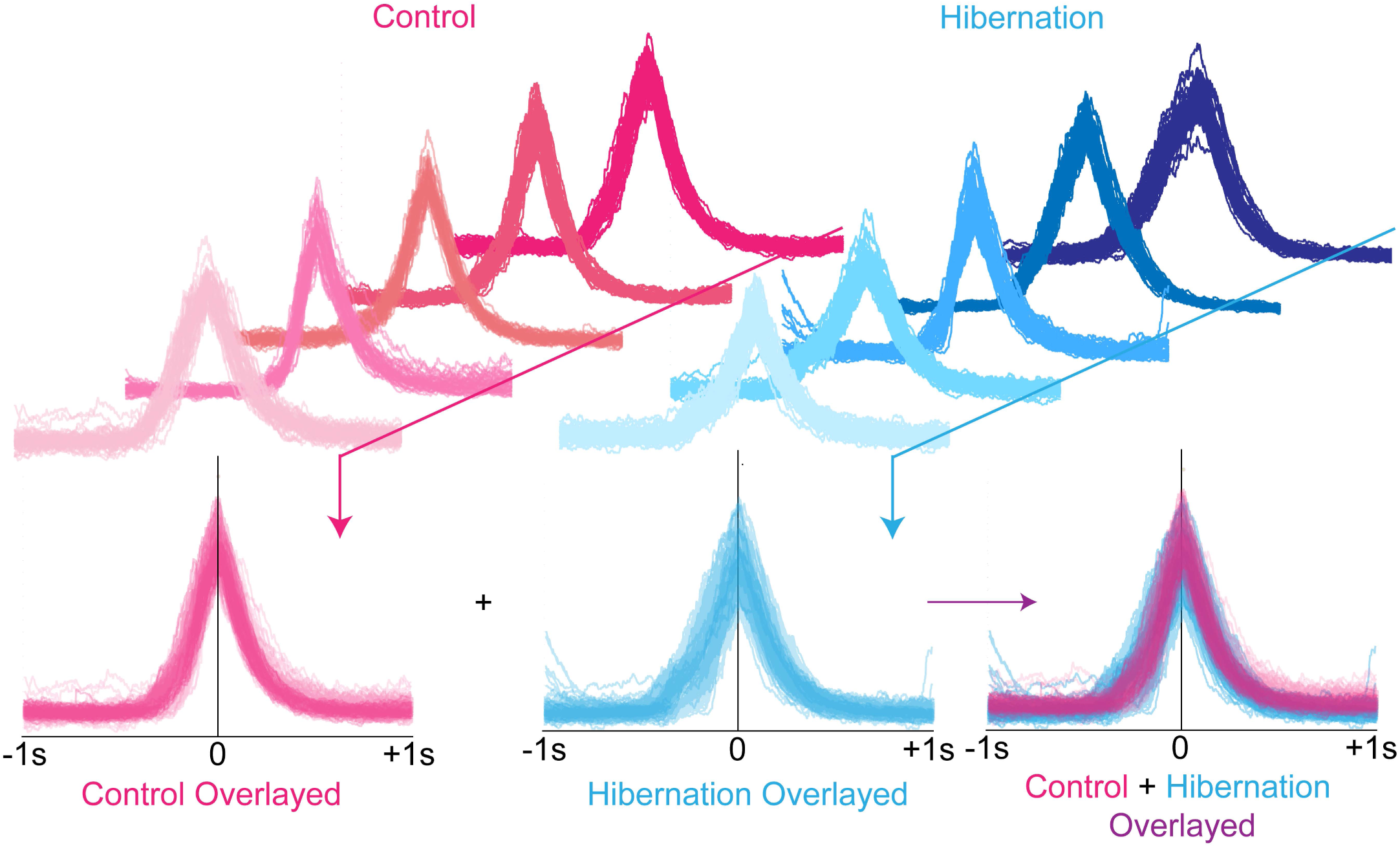
Respiratory burst morphology after hibernation closely matches controls at room temperature. Integrated vagus motor bursts (CN X) in preparations from 5 controls (red) and 5 hibernators (blue). Overlayed bursts (n=50) from individual preparations are represented in different shades from top to bottom. Overlaying all bursts from all animals from both groups demonstrates a close matching between controls and hibernators, indicating the loss of GABA_A_ receptors has no obvious impact on motor burst morphology after hibernation.

To address this, we use three groups of *ex vivo* brainstem preparations: controls, hibernators (reduced GABA signaling), and controls with GABA_A_Rs experimentally reduced with a subsaturating dose of bicuculline (2 μM). We hypothesized that control preparations with impaired GABA_A_R signaling would have similar responses to cooling that were like hibernators if GABA_A_R signaling is part of the process by which cold temperatures depress activity of the respiratory network. That is, less GABA_A_R signaling would cause enhanced activity at colder temperatures. First, we addressed the respiratory frequency. Cooling control brainstems from 20°C to 8°C expectedly reduced the respiratory burst frequency (Figure 6A). Interestingly, the decline in hibernators was less pronounced, with a temperature that produces 50% of baseline burst rate shifting to colder temperatures (Figure 6B). Decreasing GABA_A_R signaling in controls via subsaturating block of GABA_A_Rs also produced greater burst frequency at cold temperatures; however, this was more dramatic than hibernators, resulting in some networks with activity at 8°C (Figure 6C). Summary statistics are shown in Figure 6D. Two-way ANOVA revealed a significant temperature, group, and temperature by group interaction effect. These results were driven by several pairwise differences caused by hibernators and controls with bicuculline. Hibernators had statistically greater normalized burst frequency than controls at cooler temperatures (16°C: p=0.0267; Holm-Sidak Multiple Comparisons test, Figure 6D blue). In addition, bicuculline application to controls had significantly greater normalized burst frequency at lower temperatures (Figure 6D gray). These results are also summarized as the temperature at which burst frequency decreased by 50% (T_50_) (Fig 6E). Indeed, hibernators had lower T_50_ than controls, and controls with bicuculline had the lowest T_50_ values. Taken together, the network is more active at cooler temperatures when GABA_A_R signaling is reduced naturally by hibernation or by experimentally blocking GABA_A_Rs.

**Fig. 6.**
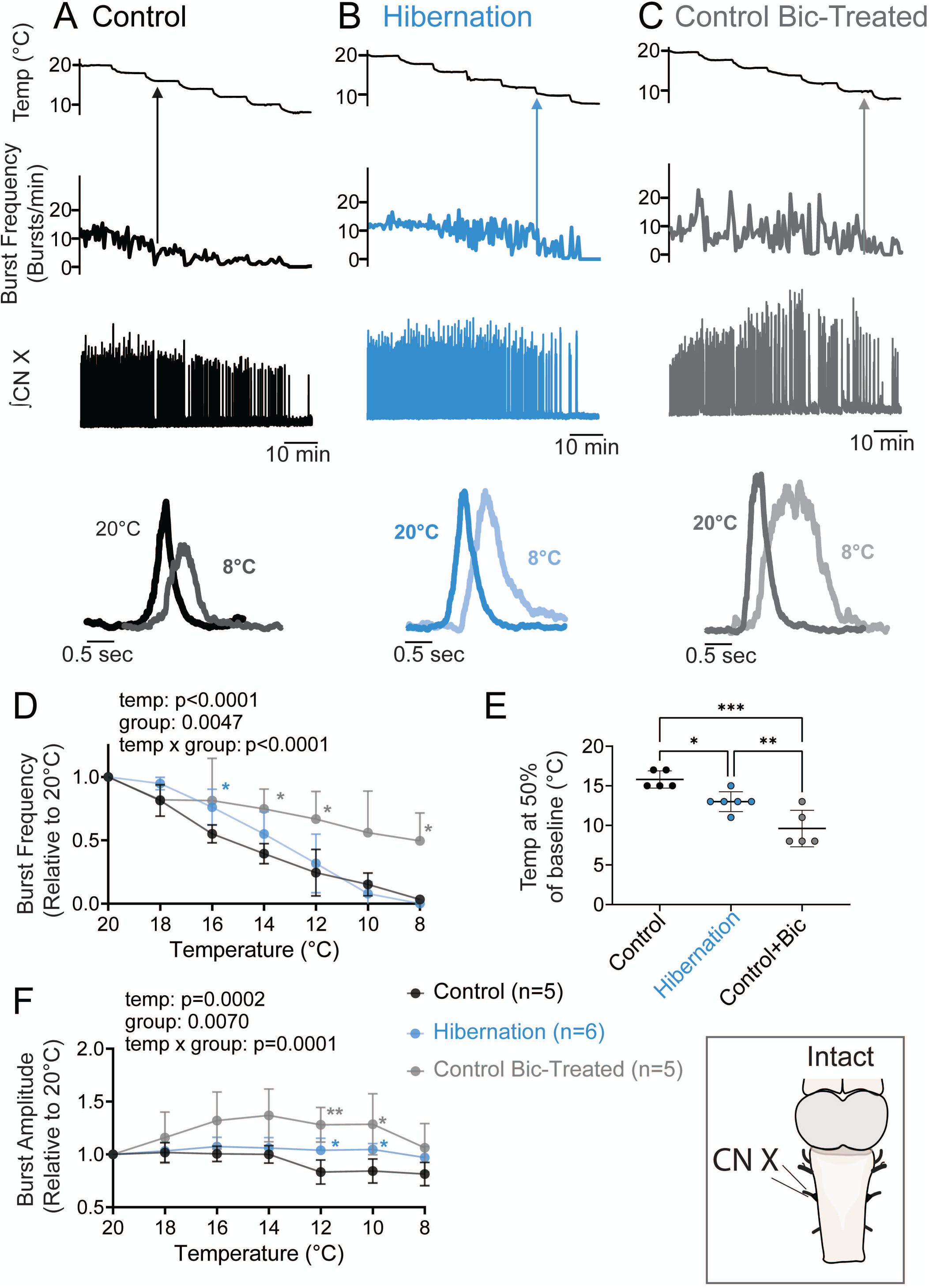
Lowering GABA_A_R signaling enhances motor performance of breathing at cold temperatures. Integrated vagus nerve activity (CN X) from intact brainstem preparations (bottom right box) showing the response of control (A, black), hibernators (B, blue), and controls treated with 2 μM bicuculline (C, gray) during cooling ramps from 20°C to 8°C. Zoomed-in traces below the compressed recordings illustrate the effects of cooling on burst amplitude. Arrows indicate the temperature at 50% of baseline burst frequency. (D) Mean data showing average temperature responses of normalized burst frequency among groups (two-way ANOVA interaction of group x temperature, p<0.0001). (E) Mean data showing a significant difference in the temperature at 50% of baseline burst rate between control, hibernators, and controls+bicuculline (one-way ANOVA; p<0.0001) (F) Mean data showing change in burst amplitude from baseline in control and hibernators during temperature changes. Controls undergo a decrease in burst amplitude during cooling, while hibernators and controls+bicuculline maintain burst amplitude throughout the cooling ramp (two-way ANOVA, p=0.0001 group x temperature interaction); * represents p<0.05, **<0.01, ***<0.001 in Holm-Sidak multiple comparisons test following one or two-way ANOVAs.

In addition to respiratory frequency, we addressed the amplitude of the motor output, as this variable relates to activation of the respiratory muscles. Like frequency, cooling also depressed motor amplitude in controls, reflecting lowered motoneuron firing or recruitment during the fictive breath (Figure 6A). In hibernators and controls in the presence of bicuculline, burst amplitude appeared to be maintained across the full temperature range (Figure 6B-C). Summary statistics are shown in Figure 6F. Two-way ANOVA revealed a significant temperature, group, and temperature by group interaction effect. These results were driven by several pairwise differences in hibernators and controls with impaired GABA_A_ signaling. Hibernators and controls with bicuculline had statistically greater burst amplitude than controls at 12°C and 10°C (12°C: hibernator vs. control: p=0.0276; bic vs. control: p=0.0039, 10°C: hibernator vs. control: p=0.0254, bic vs. control: p=0.0254) Holm-Sidak Multiple Comparisons test). Overall, while network output at warm temperatures were similar between controls and hibernators with reduced GABA signaling, these results demonstrate that the downregulation of GABA_A_R signaling by hibernation drives more robust respiratory motor activity (frequency and motor amplitude) at cooler temperatures.

## Discussion

Neural circuits face constant disturbances from the environment, placing a strong pressure on plasticity mechanisms to regulate activity. Here, we corroborate previous work suggesting GABA_A_Rs play a role in generating the respiratory motor pattern in adult frogs and identified that they also play a distinct modulatory role to depress activity during cooling. We found that the state of hibernation triggers a downregulation of synaptic inhibition which would have been expected to cause general effects on network activity. Instead, activity of the respiratory network became more resistant to decreases in activity caused by cooling in a way that would help stabilize respiratory output as the brainstem passes through a large temperature range during emergence from the hibernation environment. Therefore, compensatory forms of synaptic plasticity can expand the operating range of circuits in challenging environments without obvious impacts on activity in unstressed conditions.

### Reduced GABAergic inhibition at rhythm generating and premotor loci

Given that the respiratory motor system undergoes a massive activity challenge (inactivity during winter [15]), mechanisms consistent with activity-dependent compensation have been predicted to play a role in maintaining network integrity to promote motor behavior in the spring [16, 17, 35]. In addition to upregulation of excitatory synaptic strength on motoneurons, here we found that hibernation strongly reduced GABA_A_R signaling throughout the respiratory motor network.

The present data lead us to suggest GABA_A_R plasticity at two loci within the network (Fig. 7; top panel labeled “warm”): rhythm generating circuits and motoneurons. First, expression of the respiratory rhythm in this species involves GABA_A_Rs [24], likely through mechanisms involving reciprocal inhibition. GABA_A_Rs in premotor rhythm-generating circuits must have a diminished contribution after hibernation because motor frequency in hibernators was strongly resistant to depression caused by GABA_A_R antagonists or agonists. Although counterintuitive, these opposing pharmacological manipulations (activating and inhibiting the receptors) both lead to suppression of the rhythm. This is likely to occur because too much inhibition depresses activity by hyperpolarizing neurons, and too little prevents phasic inhibition required for rhythmogenesis. This is reminiscent of what occurs in other species where rhythm generation depends on synaptic inhibition [32, 33]. Second, activity of the respiratory rhythm generator is transmitted through excitatory and inhibitory interneuronal pathways to recruit motoneurons that drive breathing. In contrast to inhibition’s presumed role in facilitating rhythm generation, inhibition onto motoneurons has a conventional role of constraining the firing rate during respiratory-related bursting, which is reduced after hibernation (Fig. 4). As mIPSCs carried by GABA_A_Rs on motoneurons were unchanged (Fig. 3), differences in presynaptic properties of inhibitory interneurons within the respiratory network likely account for weakened GABAergic control of firing (Fig. 4).

**Fig. 7.**
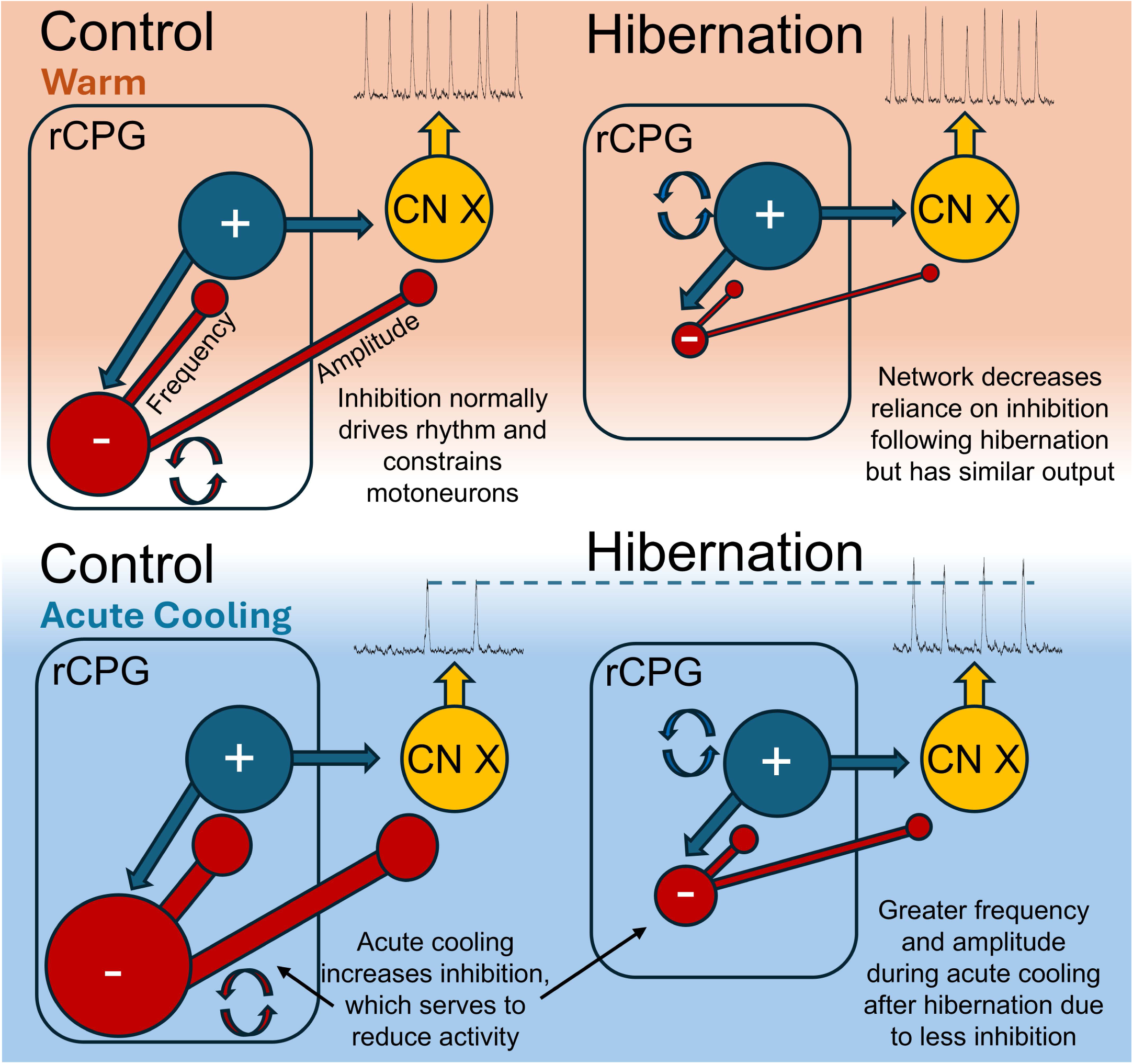
Schematic illustrating how reducing inhibition may promote breathing in the cold. In controls at warm temperatures (orange left), inhibition (red, minus symbol) plays an important role in generating the respiratory rhythm (circular arrows) and controlling the excitability to motoneurons. Following hibernation (orange right), the respiratory network decreases reliance on inhibition and potentially increases reliance on mechanisms that involve excitation (dark blue, plus symbol) for respiratory rhythmogenesis (circular arrows). Nonetheless, there is no discernable change in network motor output at warm temperatures following hibernation compared to controls. During acute cooling, inhibitory neurotransmission also plays a modulatory role and is increased. As a result, controls (blue left) have marked reduction in respiratory frequency and motor output at colder temperatures due to elevated GABAergic inhibition. In contrast, output following hibernation is resistant to depressive effects of the cold, likely with a contribution resulting from the decreased reliance on inhibition that normally slows activity and depresses motor output. Thus, output following hibernation is more robust at cooler temperatures, in part, because inhibitory neurotransmission is decreased in rhythm-generating and interneuronal pathways.

Mechanistically, it is difficult at present to parse out the variables that account for the apparent loss of network-driven presynaptic GABA (*e.g*., reduced vesicle release probability, decreased excitability of GABAergic interneurons, *etc*.), as respiratory-related synapses and GABAergic interneurons have not yet been identified in this network. Of note, rhythm and pattern generation are thought to arise, in part, from distinct populations within the respiratory central pattern generator of mammals [40]. As such, the differential effects we observed involving loss of inhibition in the respiratory circuit may stem from distinct populations within the central pattern generator itself. Overall, the specific mechanisms and network interactions await further experimentation, but the sum of our data points to the idea that rhythm-generating circuits and premotor neurons rely significantly less on GABA_A_Rs after hibernation.

Given the profound loss of GABA_A_R signaling, which is normally required to express the respiratory rhythm, one implication of these results is that some other mechanism must take over to produce breathing after hibernation. These results align well with recent work demonstrating circuits that recently recovered from an activity challenge use a different profile of ion channels to generate similar network output under baseline conditions [20, 21]. This leads us to speculate that plasticity resulting from hibernation may switch the mechanism of rhythmogenesis from one which relies heavily on inhibition to others that use excitatory synapses or intrinsic pacemaker mechanisms [24, 41] (Fig. 7; warm panel highlight shift in rhythm generating mechanisms, circular arrows). Consistent with this idea, the amphibian respiratory network appears to use voltage-dependent pacemaker mechanisms early in development, which then is replaced by network-dependent mechanisms *via* inhibition following metamorphosis [24]. Therefore, hibernation may revert the mechanisms of rhythm generation to that seen in an earlier life stage. Indeed, other aspects of breathing such as modulation of frequency by respiratory gases also revert to a juvenile-like state following hibernation [42], consistent with the idea that emerging from aquatic hibernation may share similar environmental pressures as developing air-breathing during the shift from aquatic to terrestrial habitats [43].

### Reduced GABA_A_R signaling increases the network operating range in the cold

The most intriguing aspect of these results was that networks operating with less GABA_A_R signaling after hibernation seemed to perform better at cooler temperatures (Figure 6), despite the fact that activity was largely unchanged at warm temperatures (Figure 1&5). Our data indicate that GABA_A_R signaling is part of the process by which cold normally depresses the frequency and amplitude of the motor output in control animals (Fig. 7; bottom panel labeled “acute cooling”). Although temperature effects on neural circuits are multifaceted, we draw this conclusion because subsaturating block of GABA_A_Rs in control animals opposed the typical depressive response to cooling. The mechanisms by which GABA _A_R signaling normally decreases activity in the cold seems to occur through two different processes. First, at the motoneuron level, cold may facilitate inhibitory GABAergic input that we showed to exist on these neurons (Figure. 4) to depress the population motoneuron output (Figure 6). Thus, losing this inhibition during hibernation likely explains maintenance of the burst amplitudes across the full temperature range during acute cooling (Fig. 7, bottom right, less inhibitory input in the cold). Second, our pharmacological data suggest that GABA_A_R signaling plays a role in slowing breathing frequency during cooling. Therefore, the loss of this inhibition in hibernators accounts for the ability to produce stable rhythmic activity at cooler temperatures (Fig. 6E). The network-level mechanisms remain to be uncovered; however, cold temperatures activate noradrenergic neurons of the locus coeruleus [44, 45], which then depress breathing frequency through α-adrenoreceptors [39]. Since noradrenergic signaling depresses respiratory activity mainly through GABAergic pathways [38], modulatory mechanisms involving norepinephrine may be activated by cooling to recruit GABAergic interneurons that depress the breathing frequency at cold temperatures.

Regardless of the specific mechanism by which cold temperature and GABA_A_Rs interact, reduced GABA_A_R signaling in hibernators, therefore, serves to boost respiratory frequency and motor drive at cooler temperatures, with likely relevance for restarting breathing upon exit from hibernation as the animal passes through a range of cool temperatures (Fig. 7; bottom panel labeled “cold”). As such, we interpret plasticity of GABA_A_R signaling here as “homeostatic,” as it is a change in network function that opposes the activity disturbance and enhances network excitability in an environment that otherwise depresses neuronal output. The loss of GABA_A_R signaling after hibernation seems to explain most of the maintenance of motor amplitude across temperatures, since hibernators and controls with experimentally reduced GABA_A_R signaling had similar profiles across temperatures. However, despite hibernators having higher activity at lower temperatures than naïve controls, activity still dropped to a lower extent than we observed in controls with GABA_A_R signaling experimentally reduced (Fig. 6A-D). We speculate the quantitative differences between these two conditions might point to additional plasticity mechanisms that tamp down enhanced excitability at cold temperatures to prevent ectopic breaths while submerged in cold water. While the mechanisms that serve this purpose are not fully known, we have shown that sensory input from CO_2_/pH chemoreceptors that normally stimulate breathing is reduced in hibernators [35, 42], suggesting that hibernation may also reduce excitability in some parts of the network to help keep breathing off until it is needed. The exact “switch” that restarts breathing also remains unclear, but nevertheless, our results suggest that the loss of GABA_A_R signaling prepares the network to produce strong motor bursting at cool temperatures when it does restart, which would likely contribute to restoring breathing as the animal passes through a range of colder temperature before returning to a warmer body temperature after emergence.

### Conclusions: What does it mean for plasticity to be “homeostatic?”

Since its inception [6], compensatory mechanisms within the theme of homeostatic regulation are most often portrayed as a conceptually simple (but mechanistically complex) counterbalancing of altered excitability. Our results provide evidence that mechanisms consistent with homeostatic plasticity can increase the operating range of circuits in harsh environments that otherwise disrupt activity. We acknowledge that a strict definition of “homeostatic plasticity” might not encompass the plasticity we observed. Through a homeostatic lens, one might expect network activity that drives lung ventilation to restart as compensation enhances network excitability.

However, breathing does not restart during hibernation [15], which in itself seems important for animal survival as restarting breathing prematurely could be detrimental underwater. In addition, the integrated output of the entire respiratory network does not appear obviously hyperexcitable after emergence, as breathing in vivo is comparable between controls and hibernators, which could be interpreted as inconsistent with homeostatic theory [35]. While our observations deviate from traditional definitions of negative feedback homeostasis on which the field was founded [4, 6, 46], reduced GABA_A_R signaling indeed increases the strength and frequency of respiratory motor activity in cool conditions that must be passed through to reestablish life on land after hibernation, thereby widening the circuit’s operating range across temperatures.

Intriguingly, hibernation also leads to enhanced desensitization and reduced Ca^2+^ permeability of NMDA receptors that serves to constrain excitability of this network during severe hypoxia that occurs as the animal emerges, with no impact on baseline activity of the network [47]. Therefore, multiple forms of synaptic plasticity appear to be masked at baseline, but ultimately increase the operating range for motor activity under the variety of stressors that occur during emergence from the hibernation environment. Overall, these results support the view that circuits may implement compensatory mechanisms in ways that enhance activity under specific sets of environmental constraints. These findings may have important implications for diverse neural systems, as the nervous system of most animals operate homeostatically over a range of abiotic conditions, such as changes in local tissue temperature [48], pH [49, 50], oxygen levels [47], hormonal state, and other factors that have the potential to disrupt network activity on acute timescales. By linking plasticity to a challenging environment in vivo, our results lead us to emphasize that frameworks seeking to link compensation to behavior must ultimately incorporate how plasticity is integrated over the range of environments that neural circuits may encounter. Understanding how the nervous system is tuned to survive extreme conditions in intact animals will continue to provide new insights along this path.

## Methods

### Animals

All experiments performed were approved by the Animal Care and Use Committee (ACUC) at the University of Missouri (protocol #39264). Adult Female American Bullfrogs (∼100 g weight, n= 62), *Lithobates catesbeianus*, were purchased from Rana Ranch (Twin Falls, Idaho) and were housed in 20-gallon plastic tanks containing dechlorinated water at room temperature bubbled with room air. Control frogs were acclimated for 1 week following arrival before experiments. Frogs had access to wet and dry areas. Frogs were maintained on a 12-hour light/dark cycle and fed once per week. Water was cleaned daily and changed weekly. Hibernated frogs were kept in plastic tanks under the same conditions (exception of food being withheld) as control frogs for > 1 week before temperature was gradually lowered to 4°C over 10 days in a walk-in temperature-controlled environmental chamber. Once water temperature reached 4°C, air access was blocked using a plastic screen placed in the tank. After 4 weeks of submergence, experiments commenced.

### Drugs

Strychnine hydrochloride was from Sigma-Aldrich (St Louis, MO, USA). (-)-Bicuculline methiodide, tetrodotoxin citrate, DNQX disodium salt, and Muscimol were from Hello Bio (Princeton, NJ, USA).

### Brainstem-spinal cord preparation

Brainstem-spinal cord preparations were generated as previously described [16]. Briefly, frogs were deeply anesthetized with isoflurane until visible respirations had ceased and response to foot pinch was absent. Frogs were then decapitated with a guillotine. The head was submerged in ice-cold bullfrog artificial cerebrospinal fluid (aCSF; concentrations in [mM]: 104 NaCl, 4 KCl, 1.4 MgCl_2_, 7.5 glucose, 40 NaHCO_3_, 2.5 CaCl_2_ and 1 NaH_2_PO_4_, and bubbled with 98.5% O_2_, 1.5% CO_2_; pH = 7.85) and the forebrain was pithed. The brainstem-spinal cord was then carefully removed, keeping nerve roots intact. Following the dissection, the brainstem was transferred to a chamber superfused with oxygenated aCSF. In a subset of experiments where bicuculline was bath applied to the *in vitro* preparation (described below), brainstems were transected at the spinomedullary junction before recording to avoid the contribution of spinal motor populations to non-respiratory motor activity induced through disinhibition [51]. Cranial nerve X (vagus) activity was recorded using glass suction electrodes immediately following dissection. Recordings were AC amplified (1000x, A–M Systems Model 1700, A–M Systems, Carlsborg, WA, USA), filtered (10 Hz – 5 kHz), and digitized (Powerlab 8/35 ADInstruments, Colorado Springs, CO, USA). Nerve activity was rectified and integrated (time constant =100 ms) online to allow for visualization of nerve output pattern.

Vagus nerve output was then allowed to stabilize for ≥ 1 hour before drug application. In a subset of experiments (n=10 bullfrogs, 5 per group), various does of bicuculline (500 nM, 1 µM, 5 µM, 10 µM), a GABA_A_ receptor antagonist, were superperfused for 10 minutes in order of increasing concentration and then washed out. In a different subset of experiments (n=10 bullfrogs, 5 per group), various doses (500 nM, 1 µM, 3 µM) of muscimol, a GABA_A_ receptor agonist, were added to the perfusate. Each dose was superperfused for 10 minutes in order of increasing concentration. Bicuculline (10 µM) was added to the perfusate following muscimol to reverse muscimol’s effects serving as a control that output depression was not due to some kind of drift over time, but rather activation of GABA_A_ receptors.

For temperature experiments, preparations were allowed to recover for ∼1 hr at 22°C before exposure to the temperature ramp from 20° to 8°C. Preparations from controls (n=5) and hibernators (n=6) were cooled to each temperature for 10 minutes and then the temperature was lowered by 2°C. Following cooling, preparations were rewarmed on a continuous ramp that lasted approximately 5 min. Separate control preparations (n=5) were treated with 2 μM bicuculine for 30 minutes prior to starting the cooling ramp. 2 μM was selected as it was the highest dose where respiratory-related bursts were prominent in controls. The temperature was controlled by a Warner Instruments bipolar in-line temperature controller (model CL-100; Hamden, CT), and chamber temperature was monitored directly next to the brainstem.

### Dye labeling of vagal motor neurons

Brainstems were isolated as described above. In a different group of experiments, following isolation, brainstems were transferred to a chamber superfused with oxygenated aCSF. The 4^th^ branch of the vagal nerve root, which primarily innervates the glottal dilator [52] was then isolated and suctioned into a glass pipette (bilaterally).

Fluorescent dextran dye dissolved in PBS (tetramethylrhodamine 3000 MW lysine fixable dye; Invitrogen – Thermo Fisher, Waltham, MA, USA) was then backfilled into both pipettes and left to diffuse into the nerve root for 2 hours. This generated robust labeling of vagal motor neurons as previously described [16].

### Semi-intact brainstem-spinal cord preparation

Following dye labeling, in a subset of experiments (n=15 bullfrogs) the brainstem was embedded into agarose and sliced, similar to previously described [33] with some modifications. Briefly, partially cooled, but not solidified, agarose was pipetted into a thin layer onto a scored (with fine forceps) agar block. The brainstem was then dragged onto the agarose-covered agar block, laying horizontally dorsal side up. Agarose (∼57°C) was pipetted onto the top of the brainstem caudal to obex. The agar block was then mounted on a vibrating microtome plate (Campden Vibrating Microtome 7000smz, Campden Instruments; Lafayette, IN, USA) and covered in ice-cold bubbled aCSF. The blade was zeroed at the top of the brainstem, then slices were taken caudal to rostral, stopping shortly rostral to obex which was approximately 1/3 of the distance from the hypoglossal root to the vagal root and spanned a fraction of the vagal motor pool extent. The blade was removed after each slice and the resulting attached slice was trimmed off the brainstem with fine spring scissors. Approximately 650 μm were removed from the dorsal surface in total, providing access to labeled motor neurons for recording. The brainstem was removed from the remaining agar and transferred to a 35 mm glass bottom sylgard-coated dish with a stainless-steel mesh insert. The brainstem was then pinned dorsal side up. The dish was placed in a QE-1 platform (Warner Instruments, Holliston, MA, USA) that was mounted on a fixed stage microscope (FN1, Nikon Instruments Inc., Melville, NY, USA). Once on the rig, the preparation was superperfused with bubbled aCSF either using gravity or with a peristaltic pump (Rainin Rabbit). The trigeminal nerve (CN V) was recorded using a glass suction electrode to monitor preparation output over time. Nerve recordings were AC amplified (1000x, A–M Systems Model 1700, A–M Systems, Carlsborg, WA, USA), filtered (10 Hz – 5 kHz), and digitized (Powerlab 8/35 ADInstruments, Colorado Springs, CO, USA). The trigeminal nerve was recorded instead of the vagal nerve root because of physical constraints in the bath with the patch pipette and picospritzer pipette used in experiments described below. The trigeminal provides a sufficient surrogate for efferent respiratory motor output [29, 53], as respiratory activity of the trigeminal and vagus nerves activate near-synchronously *in vitro*. Labeled motor neurons were then located and visualized using an imaging camera (Hamamatsu ORCA Flash 4.0LT sCMOS, Hamamatsu Photonics, Hamamatsu City, Japan) coupled to Nikon imaging software (NIS elements).

Whole-cell recordings were made from labeled vagal motor neurons with an Axopatch 200B amplifier (Molecular Devices) in current-clamp mode (I=0). Cells were selected based on firing pattern, that is only cells that fired spontaneous bursts of action potentials or increased firing frequency (respiratory modulated) during respiratory motor bursts were sampled, as these neurons likely innervate the glottal dilator of bullfrogs [54], which is critical to permit airflow into the lungs of bullfrogs during ventilation and as such remains inactive during hibernation. Accordingly, neurons with subthreshold respiratory input or neurons that were active only during the interburst interval were excluded from testing. Glass pipettes (∼4-7 MΩ) were filled with a solution containing (in mmol): 110 potassium gluconate, 2 MgCl_2_, 10 Hepes, 1 Na_2_-ATP, 0.1 Na_2_-GTP and 2.5 EGTA, pH ∼ 7.2 with KOH. Data were acquired in pClamp 11 software using an Axopatch 200B amplifier and Axon Digidata 1550B digitizer (Molecular Devices, San Jose, CA, USA). To determine inhibitory tone onto respiratory motor neurons, bicuculline (50 µM) was focally applied using a Picospritzer II (General Valve Corporation, Fairfield, NJ, USA) to the recorded cell following a baseline period (≥10 respiratory bursts). Bicuculline was applied using a 5 sec pulse, every 10 secs for 5-10 minutes with the picospritzer pipette slightly larger (∼2 μm diameter) than typical for focal application and placed approximately 15 μm away from the cell body. This approach was taken to saturate the cell and surrounding area with the GABA_A_R antagonist. Importantly, this approach did not recapitulate systemic effects of bicuculline on the vagal motor pool as the population spans nearly 2.5mm in the rostrocaudal axis in ranid frogs [55] and only a subset of the vagal motor pool was exposed in the preparation as described above. A washout period was used to verify changes in cell firing frequency were due to drug application and not cell drift. Most patched cells were near the surface of the tissue as the thickness of the preparation hindered cell visualization. As a result, changes in firing frequency during bicuculline application and wash were rapid, suggesting the area was saturated with drug upon administration and then quickly washed. In a subset of cells where a gigaohm seal was not formed, recordings of actional potentials were performed in the loose-patch configuration (V_hold_=0 mV; n=2 in control, and n=2 in hibernation). Regardless of recording configuration, firing rate data from recorded neurons were combined in group analysis of firing frequency, as data from loose-patch aligned with data from whole cell configuration. All voltages from current-clamp experiments were corrected for a liquid junction potential of 12 mV.

### Brain Slice electrophysiology

Following dye labeling, in a subset of experiments brainstem slices (300 µm) containing labeled motor neurons were obtained (n=11 bullfrogs). Whole-cell recordings were made from labeled motor neurons with an Axopatch 200B amplifier (Molecular Devices) in voltage clamp (V_hold_=-60 mV) mode. Glass pipettes (2.6-4 MΩ) were filled with a solution containing (in mM): 95 CsCl, 2 MgCl_2_, 10 Hepes, 1 Na_2_-ATP, 0.1 Na_2_-GTP, 10 EGTA, 1 CaCl_2_ and 10 tetraethylammonium-Cl (TEA), pH ∼ 7.2 using CsOH. Data were acquired in pClamp 11 software using an Axopatch 200B amplifier and Axon Digidata 1550B digitizer (Molecular Devices, San Jose, CA, USA). To examine miniature inhibitory postsynaptic currents (mIPSCs), tetrodotoxin (TTX; 250 nM) and DNQX (10 µM) were bath applied for 3 min to block voltage-gated Na^+^ channels and AMPA receptors. Strychnine (5 µM) was then bath applied in combination with TTX and DNQX for 2 min to block glycinergic mIPSCs and isolate GABAergic mIPSCs which were recorded for 1 min. All GABAergic mIPSCs were blocked following perfusion of bicuculline (50 µM) confirming their isolation. A 10 mV step was performed at the beginning of each recording and before each solution change to monitor series resistance (R_s_; estimated from the peak of the capacitive transient) and input resistance (R_IN_; estimated from the steady-state current). R_s_ was not compensated but remained <25 MΩ for inclusion.

### Data Analysis

For experiments using the intact brainstem preparation, respiratory burst frequency was determined from integrated vagus nerve signals using the peak analysis function in LabChart 8 (ADInstruments Inc., Colorado Springs, CO, USA). Respiratory bursts were identified based on standard metrics used in the field such as ∼1 s duration [56, 57].

Burst start and stop time points were defined as 5% of the height from baseline. In experiments where muscimol was systemically applied, data were sampled from the last 3 minutes of each condition (baseline, 500 nM, 1 µM, and 3 µM). In experiments where bicuculline was systemically applied, data were sampled for 10 minutes during each condition (baseline, 500 nM, 1 µM, 5 µM, 10 µM). In systemic bicuculline experiments where non-respiratory bursting was quantified, non-respiratory were identified by their long duration and qualitatively different shape than respiratory bursts and were counted manually. Non-respiratory bursts appeared in a variety of output patterns as previously characterized *in vitro* (Reid and Milsom, 1998). Non-respiratory bursts were not quantified during systemic muscimol exposure as they were largely absent from baseline recordings and only emerged with notable frequency following disinhibition with bicuculline. For temperature experiments, burst frequency was averaged in the last 5 minutes at each temperature, and burst amplitudes were sampled in the last minute or in the case of the failure temperature, bursts were sampled directly before failure. To compare respiratory burst amplitudes in the cold, we analyzed the change from baseline at the coldest temperature the preparation could produce activity. Across control and hibernation groups this typically occurred at 8-10°C.

For experiments using the semi-intact preparation, action potentials were detected with the peak analysis function in LabChart 8 (ADInstruments Inc., Colorado Springs, CO, USA). Following detection of action potentials, a channel was created that plotted firing frequency over time. Average firing frequency per respiratory burst was determined via the mean firing frequency from neuronal burst start to stop. Average firing frequency was sampled and averaged from 10 respiratory bursts in each condition (baseline, bicuculline, and wash).

For voltage-clamp experiments in brainstem slices, average amplitude (current measurement from baseline to peak), charge transfer (integral of the mIPSC), rise time (time from baseline to peak), and frequency of GABAergic mIPSCs (mIPSC/sec) were analyzed from one minute of gap-free recording following two minutes of exposure to TTX/DNQX/Strychnine (as described above) using the peak analysis function in LabChart 8 (ADInstruments Inc., Colorado Springs, CO, USA). Events below 7.5 pA were excluded. Events were inspected manually to ensure the accurate detection of mIPSCs. For construction of the cumulative probability histogram, the first 50 mIPSCs from the minute of data obtained from each neuron were sampled.

### Statistics

Data are presented as mean ± s.d. unless otherwise stated or shown as individual points and means to highlight individual responses. When two groups of dependent samples were compared (“before-after” experiments), a two-tailed paired t-test was used. When two groups of independent samples were compared, a two-tailed unpaired t-test was used. Non-parametric versions of these tests were used if data sets were not normally distributed. In experiments with one main effect, a one-way ANOVA was used to test for the main effect and the interaction effect. One-way ANOVA was followed up with Holm-Sidak multiple comparisons test. In experiments with two main effects, a two-way ANOVA was used to test for the two main effects and the interaction effect. Two-way ANOVA was followed up with Holm-Sidak multiple comparisons test unless otherwise specified. Cumulative distributions were compared with the Kolmogorov-Smirnov test. Significance was accepted when P< 0.05. All analyses were performed using GraphPad Prism (v9.4.1, San Diego, CA, USA).

## Declarations

Ethics approval and consent to participate: Animal experiments were approved by University of Missouri Animal Care and Use Committee protocol #39264

### Consent for publication

The authors each consent to the publication of this work.

### Availability of data and materials

Data will be made available upon request.

### Competing interests

Santin is a Guest Editor of the Special Issue “Synaptic plasticity, learning, and memory” at *BMC Biology*. The authors have no other competing interests to declare.

### Funding

National Institutes of Health R01NS114514 to JS

### Authors’ contributions

Conceived research; JS, SS. Designed research; JS, SS. Performed experiments; SS. Analyzed data; SS, JS. Drafted original manuscript; SS, JS. Edited, revised, and finalized manuscript; SS, JS

## Acknowledgements

The authors would like to thank Dr. Joe Viteri for comments on a previous version of this manuscript.

## Notes

### Competing Interest Statement

The authors have declared no competing interest.

### Summary of Updates

The revision reflects the addition of experiments that identified the functional implications of reduced GABAa signaling after hibernation. We now provide evidence that the main impact of reducing GABAa signaling is to improve network activity cooler temperatures below baseline. As we presented in the original submission, this occur without obvious changes to the network activity at baseline temperatures. These added experiments led to a major revision of the paper, placing these results in the context of the how plasticity of GABAergic inhibition can lead to different impacts on network function under different environmental conditions.

